## Supplementary Material text and figures for "Different factors control long-term versus short-term outcomes for bacterial colonisation of a urinary catheter"

(Dated: January 20, 2025)

**Contents**

**I. Background** 2
A. The Foley catheter 2
B. Catheter-associated urinary tract infection (CAUTI) 2
C. Pathogens causing CAUTI 2
D. Bacteriuria 2
E. Bacterial colonisation of the catheter 3
F. Catheter blockage 3
G. Motility of *E. coli* 3
H. Upstream swimming and catheters 4

**II. Further model details** 4
A. Bladder 4
B. Luminal flow 5
C. Connecting the different parts of the model 6
D. Connecting the outside surface and bladder 6
E. Connecting the luminal flow and luminal surface 7
Summary of the bacterial flux calculation 8
Assumptions 8
Solution 8

**III. Numerical implementation of the model** 9
A. Outside surface 9
B. Bladder 10
C. Luminal surface 11
D. Boundary and initial conditions 11

**IV. Further results of the model** 12
A. Criticality of urine production rate: washout transition 12
B. Urine production rates in the population 12
C. Bacterial surface motility can affect steady-state bacterial density in the bladder in the high urine production rate wash-out regime 12
D. Parameter regimes for the infection timescale, and their link to patient characteristics 14
E. Comparing bacterial density patterns with experimental data 15
F. Global sensitivity analysis 15

**References** 16

### I. BACKGROUND

#### A. The Foley catheter

A urinary catheter is a tube that is used to drain urine from the bladder into a drainage bag. Catheterisation can be intermittent (the catheter is removed immediately after drainage), but our study focuses on longer-term catheterisation, where the catheter is indwelling (the catheter remains in the bladder). An indwelling catheter is inserted into the bladder through the urethra, or through a hole in the abdomen (suprapubic catheter). Here we focus on the urethral case. This can occur in a hospital setting, commonly during and after surgery, or in a long-term care setting. Catheterisation is extremely common: over 90,000 people live with an indwelling catheter in the United Kingdom[1], and 30 million urinary catheters are used annually in the USA[2].

In this study, we focus on the indwelling Foley catheter. This type of catheter is a flexible tube that is inserted into the bladder via the urethra. The tube contains two lumens. Sterile water is injected into one of the lumens after insertion, to inflate a balloon just below the catheter tip. This balloon holds the catheter in place. The other lumen is used to drain urine. It has holes (eyelets) close to the catheter tip through which urine passes from the bladder, before flowing through the lumen into a drainage bag outside the body. Typical flow rates for urine passing through the catheter are 1 mL min$^{-1}$[3]. Catheter lengths range from 40 mm for women up to 160 mm, or greater, for men, with the balloon having a volume of 10 mL[3]. Modern catheters are typically made of latex or silicone, with external cross-sectional diameters of 4.0–5.3 mm (catheters are generally sized in French gauge, where 3 Fr = 1 mm; typical catheters are sized between 12 and 16 Fr)[3]. The Foley catheter remains close to its original 1930s design, although a closed drainage system (to reduce contamination from the drainage bag) was successfully introduced in the 1960s[4,5].

#### B. Catheter-associated urinary tract infection (CAUTI)

Urinary catheters are, unfortunately, prone to colonisation by bacteria. When this leads to symptoms such as fever, pain or inflammation, it is known as a catheter associated urinary tract infection (CAUTI). CAUTI are a regular part of life for patients with long-term indwelling catheters, with some studies finding incidence rates of symptomatic episodes as high as 1.1 per 100 catheterised patient-days[6]. CAUTI is also prevalent in hospital settings: in fact, CAUTI accounts for up to 40% of hospital acquired infections[2,6–11]. CAUTI incurs huge economic costs; for example, it is estimated to cost the United Kingdom £1.0 - £2.5 billion annually[3]. If bacterial colonisation of the bladder occurs without clinical symptoms, it is known as asymptomatic bacteriuria (see below).

Despite the prevalence of CAUTI, there is still limited understanding of the role of different factors in the development of infection[6,7,12–14]. Understanding these factors and the pathways to bacterial colonisation of urinary catheters may be a key step in reducing the impact of CAUTI[15].

Risk factors for CAUTI include the duration of catheterisation[6,12,16]; sex, with prevalence of CAUTI significantly higher in female hospital patients than males[16,17]; and dehydration[5,18].

#### C. Pathogens causing CAUTI

The bacterial pathogens most commonly associated with CAUTI overlap substantially with those that cause urinary tract infections more generally. With or without a catheter, infections are primarily associated with uropathogenic *Escherichia coli* (UPEC), which is present in 75% of uncatheterised urinary tract infections[19], and in 40–70% of CAUTI[7]. Other bacteria commonly isolated from CAUTI include *Klebsiella* spp, *Enterococcus* spp, *Proteus mirabilis*, and *Pseudomonas aeruginosa. P. mirabilis* has been the focus of attention in the context of CAUTI because of its tendency to form crystalline biofilms in the catheter lumen (see below, under 'catheter blockage'). CAUTI can also be caused by yeasts of the genus *Candida*[7].

#### D. Bacteriuria

It is important to clarify the difference between CAUTI and asymptomatic bacteriuria. CAUTI is defined as a bacterial infection accompanied by symptoms (inflammation, fever, pain etc), whereas bacteriuria is the presence of bacteria within urine, which can occur without symptoms[6]. Bacteriuria is extremely prevalent, occurring at an incidence rate of 3-7% per catheter day[12], and hence long-term catheterisation almost always leads to bacteriuria[20]. Bacteriuria does not necessarily need to be treated, but is often erroneously treated as CAUTI[21]. Our model does not include the response of the human host, therefore it cannot predict the presence or absence of clinical symptoms. Thus our study can predict the occurrence of bacteriuria but it cannot distinguish between CAUTI and asymptomatic bacteriuria.

##### E. Bacterial colonisation of the catheter

A number of different factors contribute to the vulnerability of catheterised patients to CAUTI infections. The catheter surface provides a direct pathway for bacteria to enter the body, harbouring biofilms and providing access to the bladder. The catheter surface is particularly vulnerable to contamination at the meatus, potentially by gut bacteria. Moreover, the presence of a foreign object in the urethra and bladder can cause trauma to the tissues there, which in turn increases their susceptibility to infection.[3,5,7,22]. The port connecting the catheter to the drainage bag also constitutes a potential point of vulnerability to infection. Although the port is theoretically sterile, the need to regularly replace the drainage bag exposes it to possible contamination, particularly for catheterised patients in the community (i.e. outside hospitals).

Catheterisation also changes the characteristics of the bladder as a site of infection. Even in the absence of a catheter, the bladder is known to retain a small volume of urine[23]; this is a rich growth medium[5] that can support high bacterial densities: up to $10^8$ CFU $g^{-1}$ (CFU = colony forming units; an experimental estimate for the number of viable cells in a sample), with doubling times as fast as 30 mins for *E. coli*[24]. With indwelling catherisation, the volume of this residual sump is increased due to a combination of factors including the location of the catheter balloon and inlets, the cessation of tidal drainage in favour of continuous drainage, and hydrostatic pressure differentials, particularly as a result of kinks in the catheter tubing[25]. The prevention of tidal drainage also disrupts the self-cleaning nature of the bladder and urethra, preventing the regular flushing out of bacteria[2,3]. Thus, the catheter can act as a reservoir for bacteria to spread into the bladder, but so too can the residual urine within the bladder act as a reservoir from which the catheter can become infected. It is likely that the epithelial cells that line the bladder walls also have a key role to play in infections. Some patients experience recurrent urinary tract infections due to persistent intracellular bacterial communities in the epithelial cells[26]; the same mechanism could also cause recurrent CAUTI[27].

It has been established that around two thirds of catheter-associated infections originate from bacterial growth on the outside of the catheter[27]. Most of the remaining infections are attributed to bacteria infecting the drainage bag or port and then ascending the inside (luminal) surface of the catheter. Only around 5% of infections are due to contamination at the point of catheter insertion[7].

##### F. Catheter blockage

CAUTI is not merely an unpleasant inconvenience for sufferers; it also brings risk of serious consequences, including kidney infections, bloodstream infections and tissue damage within the bladder[28]. A common consequence of CAUTI is blockage of the catheter, which will, at best, result in urine bypassing the catheter, and at worst can lead to the backflow of infected urine into the kidneys[19]. Catheter blockage is sometimes caused by the formation of a thick biofilm by bacteria such as *P. aeruginosa*[29], however more commonly the cause is crystalline biofilms formed by *P. mirabilis*[27,30]. *P. mirabilis* hydrolyses urea, producing ammonia, which increases the alkalininty of the urine, leading to precipitation of crystals of struvite and apatite[27,30].

##### G. Motility of *E. coli*

The motility of bacterial cells plays a key role in colonisation of urinary catheters. For *E. coli*, motility has been extensively characterised in *in vitro* experiments. *E. coli* cells swim in liquid media by rotating their (helical) flagella together[31], generating an anisotropic frictional force, and propelling the cells forwards[32]. *E. coli* has a characteristic 'run-and-tumble' style of motion, as cells periodically reverse the direction of some of their flagella, disrupting the rotation, and reorienting the cell to swim in a different direction[33]. At long length- and time-scales, this run-and-tumble motion can be mapped onto the statistical physics model of a random walker[31], characterised by an active diffusion coefficient – $D \sim 100\ \mu m^2 s^{-1}$[34]. The motility of *E. coli* on or near surfaces is more complicated, and in general is less well-characterised.

Our decision to describe the bacterial surface motility in our model by a diffusive term enables the parameterisation of the motility by a single experimentally determinable parameter: an effective diffusion coefficient $D_S$. However, it carries a significant limitation, since it reduces the complexity of bacterial motility on the surface to just diffusion and hence overlooks any more complicated behaviours, such as swarming (swarming cells are highly motile[35], performing super-diffusion[36]).

It is important to highlight the large range in literature estimates for the bacterial surface diffusivity. This arises in part from difficulties in experimentally assessing bacterial motility on catheter surfaces: bacterial motility on the catheter surface depends significantly on surface 'wetness'[35] (that is, bacteria typically exhibit slow 'twitching' styles of motility on dry surfaces, whereas cells can often more effectively utilise their flagella – and hence move faster – when there is a fluid layer covering the surface), and the surface properties of the catheter differ significantly *in vitro* versus *in vivo*[37], due at least in part to host-catheter interactions that result in the formation of a conditioning film on the catheter surface[19]. However, the uncertainty is also a consequence of describing the bacterial motility as simple diffusion. In reality, there are enormous differences in motility from species to species[38], but even within a single bacterial strain there can be multiple modes of motility. For example, a population of cells growing within a biofilm often have very limited motility, resulting in a population that spreads primarily by growth[29] ($D_S \sim 10^{-9}$ mm$^2$s$^{-1}$), while bacterial cells actively crawling or swimming over a surface show motility that is heavily influenced by surface 'wetness'[34,35,39] (from $D_S \sim 10^{-8}$ mm$^2$s$^{-1}$ to $D_S \sim 10^{-4}$ mm$^2$s$^{-1}$). This large range of plausible values for the bacterial surface diffusivity results in uncertainty in predictions of infection timescales from our model.

##### H. Upstream swimming and catheters

The hydrodynamics of a micro-swimmer (such as a bacterial cell) can be calculated close to a surface (such as a catheter), and often differs significantly from the far-field hydrodynamics[40]. In shear flow close to a surface, for low-to-moderate shear rates, *E. coli* consistently swims upstream; this is due to the local shear rate causing cells to pivot and orient to point into the flow[41]. This phenomenon of upstream swimming has previously been discussed in the context of urinary catheters, with the suggestion that *E. coli* might ascend from a contaminated drainage bag to the bladder in $\sim 5$ hrs[42].

However, in clinical settings, bacteriuria occurs over a timescale of days rather than hours (around $\sim 25\%$ of patients are bacteriuric within 2 weeks[27]). This suggests that upstream swimming may not be the dominant means by which bacteria ascend catheters. In fact, bacterial contamination of the drainage bag is a comparatively rare event: the majority of catheter-associated infections originate instead from bacterial growth on the outside surface of the catheter. In addition, bacteria readily adhere to the catheter surface, since uropathogenic strains of *E. coli* express many more adhesins than laboratory strains[27], and the catheter surface rapidly develops a conditioning layer of proteins such as fibrinogen following catheterisation[19,43].

Moreover, if we calculate the shear rate of the urine flow at the catheter surface for the parameter values assumed in Table 1 of the main text, we find the shear rate to be 21 s$^{-1}$, well above the critical threshold for upstream swimming to occur ($\lesssim 10$ s$^{-1}$)[41]. It seems unlikely therefore that the phenomenon of upstream swimming is significant for the bacterial colonisation of urinary catheters.

#### II. FURTHER MODEL DETAILS

##### A. Bladder

We use a logistic equation to describe growth of bacteria in the bladder (Fig. S1a). Previous work by Gordon and Riley[23] assumed exponential bacterial growth in a model for micturition dynamics and urinary tract infections. However, an exponential growth model is unbounded, rapidly diverging toward infinite bacterial densities, rendering it appropriate only for short timescales. Logistic growth represents the simplest modification that can be made to an exponential growth model to enforce an upper bound on the population density.

While the logistic growth model that we use is simple, it has no mechanistic justification. An alternative model would be that of a chemostat: a device in which bacteria grow under conditions of continuous dilution, which closely resembles the situation of bacterial growth in a catheterised bladder. However, chemostat theory describes bacterial growth only on a single, simple limiting substrate, e.g. glucose as the sole carbon source. Urine is not a simple substrate, and no chemostat model has yet been developed to describe quantitatively bacterial growth in urine. As a minimal model, therefore, assuming logistic growth dynamics may be a reasonable approximation.

As discussed in the main text, the value of the dilution rate $k_D$ (which is controlled by the urine production rate $\lambda$ and the bladder volume $V$ via $k_D = \lambda/V$) is critical for the behaviour of our model. The dilution rate governs both the steady state bacterial density in the bladder (Fig. S1b), and the timescale over which the steady state is attained (Main text Fig. 2b). This dependence of the model on the dilution rate is very similar to the behaviour of a previous (uncatheterised) micturition model, in which there is a critical relationship between the urine dilution rate and the

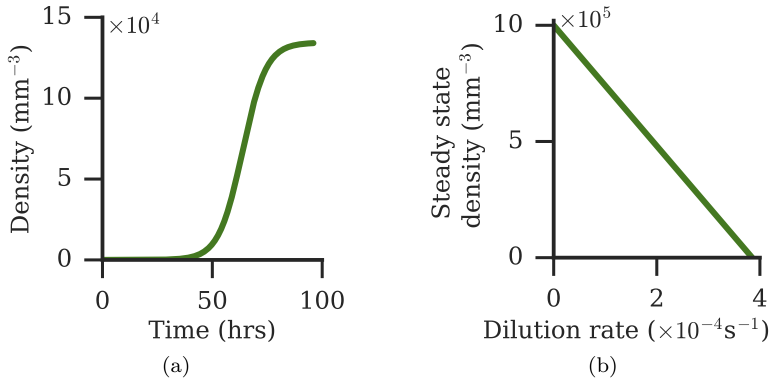


Figure S1: **Importance of bladder parameters.** (a) Behaviour of the bladder model (Eq. 2 of the main text) when uncoupled from the rest of the system: the dynamics of the bacterial density as the system approaches the steady state. Initial condition is $\rho(t=0) = 1$ mm$^{-3}$. (b) Dependence of the steady state solution of Eq. 2 on the dilution rate, $k_D$. The steady state bacterial density decreases linearly as the dilution rate increases. All parameters except the dilution rate are as in Table 1 of the main text.

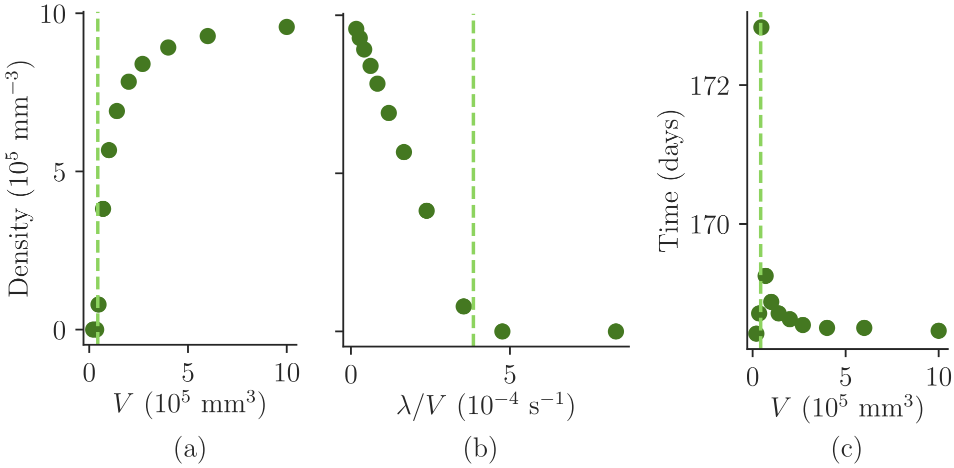


Figure S2: **The residual urine volume also determines the dilution rate.** (a) The effect of varying the volume of residual urine, $V$ on the steady state bacterial density in the bladder. The vertical line (▬) marks the washout transition, $V = \lambda/r_B$. (b) The same data points as in (a), however the x axis now plots the dilution rate, $k_D$, where each point was obtained from a simulation with fixed $\lambda$ but varying $V$. (c) The effect of varying the residual urine volume on the time taken to steady state bacterial density in the bladder. $V$ is varied within the physiologically relevant range: 10 - 1000 mL. The surface bacterial motility is $D_S = 10^{-8}$ mm$^2$ s$^{-1}$. All other parameters are as in Table 1 of the main text.

bacterial growth rate[23]. However in our model the presence of the catheter means that the bacterial density in the bladder does not go to zero in the high dilution rate ‘wash-out’ state.

Since the urine production rate is incorporated into the bladder model through the dilution rate ($k_D = \lambda/V$), varying the residual urine sump volume in the bladder has similar effects on the bladder dynamics to varying the urine production rate (Fig. S2). Comparing Fig. S2b with Fig. S1b, the same linear behaviour can be observed. Similarly, comparing Fig. S2c with the upper inset of Fig. 3a in the main text, the characteristic spike in timescales of the washout transition can be seen. We note, however, that the luminal flow dynamics are independent of the residual urine sump volume (although they do depend on the urine production rate).

##### B. Luminal flow

The hydrodynamics of flow through a pipe is well-established; the properties of the flow are determined by its Reynold’s number[44]. We can calculate the Reynold’s number for a ‘typical’ catheter of lumen radius $R = 1$ mm, and urine flow rate of $\lambda = 1$ mL min$^{-1}$:

$$\mathrm{Re} = \frac{\lambda}{\pi R \nu} = \frac{\frac{1}{60}10^{-6}}{\pi 10^{-3} \cdot 0.83 \cdot 10^{-6}} \approx 6 \tag{S1}$$

where $\nu = 0.83$ mm$^2$s$^{-1}$ is the kinematic viscosity of urine at 37 °C[45]. Thus, for urine flowing through a catheter, the Reynold's number is low, $\approx 6$, and the flow is Poiseuille (Fig. S3a).

In Eq. 3 of the main text, we assume that bacteria diffuse radially within the urine flow. We neglect both diffusion of bacteria in the longitudinal direction, and growth of bacteria within the flow. Examining the timescales of the problem justifies this. The timescale of bacterial growth is the doubling time, $\log 2/r_B = 1.8 \times 10^3$ s. The characteristic timescales for the radial and longitudinal diffusive processes are $R^2/D_B = 1 \times 10^4$ s and $L^2/D_B = 1.6 \times 10^7$ s. The typical timescale for the flow is the average time taken for urine to pass through the catheter, $\pi R^2 L/\lambda = 7.5$ s. The timescale of convective flow down the catheter is clearly much faster than the timescales of either of the diffusional processes, or growth. However we choose to include radial diffusion in the model because, although slow, it has significant effects over small radial distances, such as close to the catheter wall. This is important for calculating the bacterial flux from the luminal flow to the luminal catheter surface (Fig. S3b), which we derive below ('Connecting the luminal flow and luminal surface').

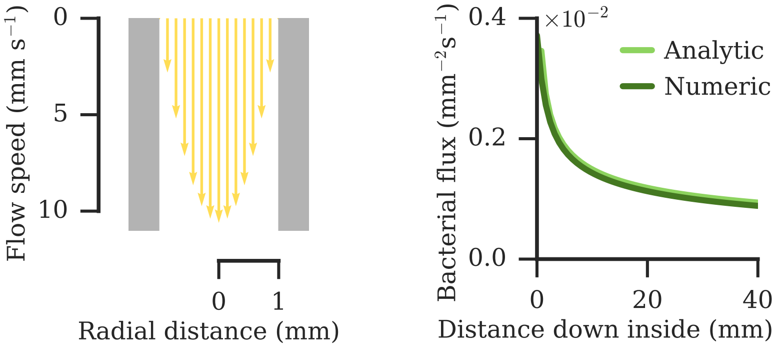


Figure S3: **Solutions for the luminal flow profile and deposition onto the luminal surface.** (a) The flow of urine through the catheter is parabolic (Poiseuille flow). (b) Bacterial flux onto the luminal surface. This is the number of bacteria sticking to the surface per unit time, for the density distribution shown in Fig. 2c of the main text. Dark green line: results of numerically simulating Eq. 3 and Eq. 10 of the main text. Light green line: the analytic result of Eq. 11 of the main text, for $x > 0.3$ mm.

#### C. Connecting the different parts of the model

The different parts of the model are connected through coupling terms which preserve the total bacterial number. The various couplings are illustrated in Fig. S4a, and described in detail in the following sections.

#### D. Connecting the outside surface and bladder

The top of the catheter is within the bladder and immersed in urine. Therefore, bacteria may detach from the outside surface of the catheter and join the planktonic population within the bladder. Conversely, bacteria within the urine in the bladder may stick to the outside catheter surface. In our model, this coupling is incorporated into the outside surface equation (Eq. 1 of the main text) as a flux at the top boundary. For the bladder, the coupling takes the form of a source term added to Eq. 2 of the main text. The coupling must conserve bacterial number, so the total number of bacteria moving to the bladder must equal the number leaving the catheter surface.

To derive the coupling terms (Eqs. 5 and 6 of the main text), we consider that only bacteria that are in the region of 'contact' between the outside surface and bladder can migrate. Therefore, we need to define a contact surface (at the top of the catheter) and a contact volume (in the bladder surrounding the catheter) within which bacteria can transfer. Fig. S4b illustrates how we define these contact geometries. Assuming that a length $l$ of catheter is exposed within the bladder, and that bacteria in the bladder within a distance $w$ have a chance of sticking, we can write down

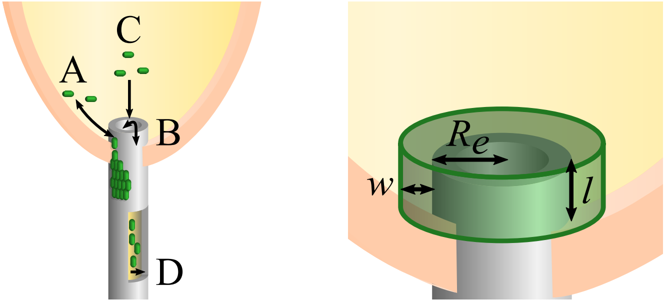


(a) (b)

Figure S4: **Connecting the different parts of the model.** (a): Overview. A: bacterial transfer between the top of the outside surface and the bladder. B: Bacterial transfer between the top of the outside surface and the top of the luminal surface. C: Bacterial transfer between the bladder and the luminal flow. D: Bacterial deposition from the luminal flow onto the luminal surface. (b) Illustrating the 'contact area' and 'contact volume'. Some bacteria within the bladder are close to the external catheter surface. At the top of the catheter, some bacteria on the surface are immersed in urine. If the catheter protrudes a distance $l$ into the bladder, then there is a total surface contact area on the catheter of $S_c = 2\pi R_e l$. We suppose that bacteria within the bladder are 'in contact' with the catheter if they are within a distance $w$, and we simplify this contact volume to be a cylinder of radius $R_e + w$ and height $l + w$. The contact volume is then $V_c = \pi(R_e + w)^2(l + w) - \pi R_e^2 l$.

the contact surface area and volume:

$$\begin{aligned} S_c &= 2\pi R_e l \\ V_c &= \pi(R_e + w)^2(l + w) - \pi R_e^2 l. \end{aligned} \tag{S2}$$

Here $R_e$ is the external catheter radius, and $l$ and $w$ are as shown in Fig. S4b.

Since we need to conserve the bacterial number, we must consider the absolute number of bacteria within the contact regions. $N_s = S_c \cdot n(L, t)$ is the number of bacteria on the catheter surface that are 'in contact' with the bladder, and $N_b = V_c \cdot \rho(t)$ is the number of bacteria in the bladder that are 'in contact' with the catheter. Therefore, the flux of bacteria to the bladder is

$$\underbrace{k_d N_s}_{\text{detaching}} - \underbrace{k_a N_b}_{\text{attaching}} \tag{S3}$$

where $k_d$ and $k_a$ are respectively the rates at which bacteria detach from and attach to the catheter surface in the presence of urine; and $N_s$ and $N_b$ are the numbers of bacteria at the boundary on the outside of the catheter and in the bladder. To set reasonable values for $k_d$ and $k_a$ we make the following assumptions: any bacteria that are born on the exposed catheter surface detach, so $k_d = r_S$; and bacteria in the bladder diffuse and stick to the surface irreversibly upon contact, so $k_a$ is the Smoluchowski diffusion rate limited constant[46]. This makes it possible to write down the coupling terms. In the bladder there is a source term for the density,

$$\frac{k_d S_c n(0, t) - k_a V_c \rho(t)}{V}, \tag{S4}$$

where $V$ is the total residual urine volume of the bladder. At the top of the catheter there is a corresponding surface density flux at the boundary:

$$\frac{k_a V_c \rho(t) - k_d S_c n(0, t)}{S_c}. \tag{S5}$$

Eqs. S4 and S5 correspond to Eqs. 5 and 6 of the main text.

#### E. Connecting the luminal flow and luminal surface

Bacteria from the luminal flow are adsorbed onto the luminal surface. To calculate this numerically requires simulating the luminal flow, which is computationally expensive as it is a 2-dimensional problem, unlike the 1-dimensional surfaces elsewhere in the model.

Instead we model the adsorption of bacteria on the luminal surface using diffusion boundary theory, established by Levich[47]. In this theory, bacteria are modelled as particles which diffuse in the flow and adsorb on contact with the surface. The flow close to the surface becomes depleted of particles due to adsorption, hence particle deposition decreases with downstream distance (see Fig. 2c of the main text). The nature of this decrease depends on the geometry of the flow[47].

By making some assumptions, in particular neglecting surface forces (the so-called 'Smoluchowski-Levich' assumption)[48], and assuming the presence of a diffusion boundary layer[47], it is possible to write an analytic approximation for this bacterial flux to the catheter surface. The calculation leading to this approximation is outlined below; it follows the path laid out by Levich[47] (in particular, chapter 2.20, 'Diffusion in laminar flow in a tube').

*Summary of the bacterial flux calculation*

In the following we show how the maximal deposition of bacteria on the surface of a pipe can be approximated by

$$j(x) = D_B \left.\frac{\partial \sigma}{\partial y}\right|_{y=0} = 0.5835 D_B \rho \sqrt[3]{\frac{\lambda}{R^3 D_B}} \frac{1}{\sqrt[3]{x}}, \tag{S6}$$

which holds provided the Reynolds number $\mathrm{Re} < 2500$ and $L \ll \lambda/D_B$. Here $j(x)$ is the bacterial flux to the surface, $D_B$ is the bacterial bulk diffusivity, $\rho$ is the initial bacterial density (at the entrance of the pipe), $\lambda$ is the volume flow rate, $R$ is the pipe radius, $x$ is the longitudinal displacement, $L$ is the pipe length, $\sigma(x, y)$ is the bacterial density within the fluid, and $y$ is the perpendicular displacement from the surface (i.e. a planar approximation of the radial co-ordinate).

We first verify that the conditions specified for Eq. S6 to be valid hold for our catheter model. Typical parameter values are $L = 40$ mm, $R = 1$ mm, $\lambda = 16.7$ mm$^3$s$^{-1}$ and $D_B = 10^{-4}$ mm$^2$s$^{-1}$ (see Table 1 of the main text), so that $\lambda/D_B \sim 10^5$ and $L \ll \lambda/D_B$. Additionally, $\mathrm{Re} = 6$, so $\mathrm{Re} < 2500$. Hence we can calculate the maximal deposition flux to be

$$j(x) \approx 0.003 \frac{\rho}{\sqrt[3]{x}}. \tag{S7}$$

Comparing the flux calculated from the numerical simulation with the analytic approximation shows good agreement, with more bacteria sticking at the top than further down (Fig. S3b).

*Assumptions*

Next, we list the assumptions that are required to derive the analytical solution.

1. All bacteria that contact the surface stick, i.e. the 'perfect sink' assumption. This is an absorbing boundary condition and leads to a maximal estimation for the deposition flux.

2. There are no external forces, except for the pressure differential driving the flow. This is the so-called 'Smoluchowski-Levich' assumption[48].

3. The flow within the pipe is laminar. This requires $\mathrm{Re} < 2500$, and is valid beyond the initial hydrodynamic inlet region, $x > h \sim R \cdot \mathrm{Re}/27$. For our model catheter, $\mathrm{Re} \sim 6$, and so $h \sim 0.2$ mm. Since the length of a catheter is 40–160 mm, the majority of the modelled catheter is in the laminar flow regime.

4. The diffusion profile is not fully established, and so there exists a thin diffusion boundary layer in which bacteria are depleted close to the surface (we note that the diffusion boundary layer becomes broader as the fluid travels down the catheter, since the diffusing bacteria spread out in the flow). The diffusion profile becomes fully established at a distance $H \sim \lambda/D_B$ down the pipe. For $x \ll H$, the boundary layer is thin, and can be approximated as planar. Therefore in relation to the catheter, we require $L \ll \lambda/D_B$, which is the regime we are working in.

*Solution*

We now explain how the solution is obtained. In the diffusion inlet region ($h < x \ll H$), diffusion occurs only close to the surface, and hence we can take a planar approximation, defining $y = R - r$ as a small variable. Our aim is then to solve the planar advection-diffusion equation,

$$v\frac{\partial \sigma}{\partial x} = D_B \frac{\partial^2 \sigma}{\partial y^2} \tag{S8}$$

where $v = \frac{2\lambda}{\pi R^2}\left(1 - \frac{r^2}{R^2}\right) = \frac{2\lambda}{\pi R^4}\left(2Ry - y^2\right)$. Since $y$ is small, we expand $v$ keeping only terms to first order in $y$, $v \approx \frac{4\lambda}{\pi R^3} y$. Then we can write

$$\frac{4\lambda}{\pi R^3} y \frac{\partial \sigma}{\partial x} = D_B \frac{\partial^2 \sigma}{\partial y^2}, \tag{S9}$$

with boundary conditions

$$\sigma = 0 \;\text{ at } y = 0, \qquad \sigma = \rho \;\text{ as } y \to \infty. \tag{S10}$$

We next introduce the dimensionless variable

$$\eta = \left(\frac{4\lambda}{\pi R^3 D_B}\right)^{\frac{1}{3}} \frac{y}{x^{1/3}}, \tag{S11}$$

to obtain

$$\frac{d^2\sigma}{d\eta^2} + \frac{1}{3}\eta^2 \frac{d\sigma}{d\eta} = 0, \tag{S12}$$

which can then be solved analytically. This gives an expression for $\sigma$,

$$\sigma(\eta) = \frac{\rho\sqrt[3]{3}}{\Gamma(\frac{1}{3})} \int_0^\eta \mathrm{e}^{-\frac{1}{9}z^3}\, dz, \tag{S13}$$

where $\Gamma$ is the Gamma function. Finally we obtain the deposition flux

$$\begin{aligned} j(x) &= D_B \left(\frac{\partial \sigma}{\partial y}\right)_{y=0} \\ &= D_B \left(\frac{d\sigma}{d\eta}\frac{\partial \eta}{\partial y}\right)_{y=0} \\ &= \frac{D_B \rho}{\Gamma(\frac{1}{3})} \sqrt[3]{\frac{12\lambda}{\pi R^3 D_B}} \frac{1}{\sqrt[3]{x}} \\ &= 0.5835 D_B \rho \sqrt[3]{\frac{\lambda}{R^3 D_B}} \frac{1}{\sqrt[3]{x}}, \end{aligned} \tag{S14}$$

where the constant prefactor 0.5835 is obtained by numerical evaluation of $\frac{1}{\Gamma(1/3)} \sqrt[3]{12/\pi}$.

#### III. NUMERICAL IMPLEMENTATION OF THE MODEL

The model is implemented as three discrete equations (outside surface, bladder, and luminal surface), coupled to one another and stepping forward in time. A forward-time centred-space (FTCS) method is used for both the outside and luminal surface, with the bladder being solved with a forward Euler method.

##### A. Outside surface

Recall Eq. 1 from the main text:

$$\frac{\partial n}{\partial t} = D_S \frac{\partial^2 n}{\partial x^2} + r_S n \left(1 - \frac{n}{\kappa_S}\right).$$

This can be discretised (with FTCS) as:

$$n_p^{i+1} = \frac{D_S \Delta t}{\Delta x^2}\left(n_{p+1}^i - 2n_p^i + n_{p-1}^i\right) + (1 + r_S \Delta t)\, n_p^i - \frac{r_S \Delta t}{\kappa_S}\left(n_p^i\right)^2, \tag{S15}$$

where $n_p^i$ is the outside bacterial surface density, $n(x,t)$, at the $p$th discrete position, and the $i$th time step; $\Delta t$ is the time step; and $\Delta x$ is the spatial discretisation. This has been shown to be numerically stable provided[49]:

$$\begin{aligned} \Delta x &< \sqrt{\frac{D_S}{r_S}} \\ \Delta t &< \frac{\Delta x^2}{2D_S} \end{aligned} \tag{S16}$$

which for the parameter values in Table 1 in the main text are 0.7 mm and 2000 s respectively. In this work, the values chosen for the step sizes were $\Delta x = \frac{1}{2}\sqrt{\frac{D_S}{r_S}}$, and $\Delta t = \frac{\Delta x^2}{20 D_S}$, which for the parameter values given in Table 1 of the main text are $\Delta x = 0.3$ mm, and $\Delta t = 45$ s.

Since bacteria are free to diffuse between the outside and luminal surfaces of the bladder, if there are $N = L/\Delta x$ discrete spatial positions, then $n_{N+1}^i = m_0^i$ (main text Eq. 7). The outside surface is coupled with the bladder, with flux given by Eq. S5:

$$\frac{k_a V_c \rho(t) - k_d S_c n(0,t)}{S_c},$$

so at the top of the catheter, $n_0$ is incremented as

$$n_0^{i+1} = \frac{D_S \Delta t}{\Delta x^2}\left(m_0^i - 2n_0^i + n_1^i\right) + (1 + r_S \Delta t)\, n_0^i - \frac{r_S \Delta t}{\kappa_S}\left(n_0^t\right)^2 + \frac{1}{2\pi R_e \Delta x}\left(k_a V_c \rho^i - k_d S_c n_0^i\right). \tag{S17}$$

##### B. Bladder

Including the coupling with the outside surface, Eq. 2 from the main text becomes:

$$\frac{d\rho}{dt} = r_B \rho \left(1 - \frac{\rho}{\kappa_B}\right) - k_D \rho + \frac{k_d S_c}{V} n(x{=}0) - \frac{k_a V_c}{V}\rho. \tag{S18}$$

Discretising this with a forward Euler method gives

$$\rho^{i+1} = \rho_i + \Delta t\left(\left(r_B - k_D - \frac{k_a V_c}{V}\right)\rho^i - \frac{r_B}{\kappa_B}\left(\rho^i\right)^2 + \frac{k_d S_c}{V} n_0^i\right), \tag{S19}$$

where $\rho^i$ is the bladder bacterial volume density at the $i$th time step. The stability can be evaluated by comparison with the logistic map, as follows. Since we know that $n_0^i$ is directly coupled to $\rho^i$, we can approximate Eq. S19, as

$$\begin{aligned} \rho^{i+1} &= \left(1 + \Delta t\left(r_B - k_D - \frac{k_a V_c}{V} + \frac{k_d S_c}{V}\right)\right)\rho^i + \frac{r_B \Delta t}{\kappa_B}\left(\rho^i\right)^2 \\ &= (1 + A\Delta t)\rho^i + \frac{r_B \Delta t}{\kappa_B}\left(\rho^i\right)^2 \end{aligned} \tag{S20}$$

which, with a change of variables $u^i = -\frac{r_B \Delta t}{\kappa_B (1 + A\Delta t)}\rho^i$, becomes

$$u^{i+1} = (1 + A\Delta t)\, u^i (1 - u^i), \tag{S21}$$

which stably converges to its non-zero equilibrium (see Murray chapter 2.3[50]) for $1 < 1 + A\Delta t < 2$, i.e.

$$\Delta t < 1 \Bigg/ \left(r_B - k_D - \frac{k_a V_c}{V} + \frac{k_d S_c}{V}\right) \sim 10^4 \text{ s}. \tag{S22}$$

##### C. Luminal surface

Recall Eq. 4 from the main text:

$$\frac{\partial m}{\partial t} = D_S \frac{\partial^2 m}{\partial x^2} + r_S m \left(1 - \frac{m}{\kappa_S}\right) + j(x).$$

The bacterial flux, $j(x)$, comes from the deposition of bacteria onto the luminal surface from the urine flow out of the bladder. Here we take the analytic approximation for the bacterial flux, as given by Eq. S6 above (also main text Eq. 11),

$$j(x) = 0.5835 D_B \rho \sqrt[3]{\frac{\lambda}{R^3 D_B}} \frac{1}{\sqrt[3]{x}}.$$

This analytic solution is valid for $h < x \ll H$, i.e. the region in which the hydrodynamic flow is established, but the diffusive boundary layer is still small. The hydrodynamic establishment distance, $h$, is the distance at which the hydrodynamic boundary layer thickness is equal to the catheter radius. From Levich[47], defining the hydrodynamic boundary layer thickness as the thickness at which the flow speed is 90% of the main flow speed, this establishment distance occurs at

$$h \sim \frac{\mathrm{Re} \cdot R}{27} \sim 0.22 \text{ mm}. \tag{S23}$$

The expected behavious at the very top of the catheter lumen, $x < h$, is highly dependent on the exact geometry of the catheter, which is not incorporated into this model. Instead, knowing that the bacterial deposition must always be finite, we take a zeroth order approximation that the flux for $x < h$ is constant, and $j(x < h) = j(h)$. Since this is only a very small region of the catheter, this approximation has little impact on the results of the model.

Thus, we can discretise the luminal surface in a manner similar to the outer surface of the catheter; using a FTCS method gives

$$m_p^{i+1} = \frac{D_S \Delta t}{\Delta x^2} \left(m_{p+1}^i - 2m_p^i + m_{p-1}^i\right) + (1 + r_S \Delta t)\, m_p^i - \frac{r_S \Delta t}{\kappa} \left(m_p^i\right)^2 + 0.5835 \left(\frac{\lambda D_B^2}{R^3}\right)^{1/3} \rho^i (p\Delta x)^{-1/3}, \tag{S24}$$

where $m_p^i$ is the luminal bacterial surface density at the $p$th spatial point, and the $i$th time step. This is numerically stable under the same conditions as the outside surface, provided the conditions discussed above hold for the validity of $j(x)$. That is, $L \ll 10^5$ mm. At the top of the catheter, bacteria can diffuse between the luminal surface and the outside surface, so at the top of the catheter $m_0$ is incremented as

$$m_0^{i+1} = \frac{D_S \Delta t}{\Delta x^2} \left(m_1^i - 2m_0^i + n_0^i\right) + (1 + r_S \Delta t)\, m_0^i - \frac{r_S \Delta t}{\kappa} \left(m_0^i\right)^2 + 0.5835 \left(\frac{\lambda D_B^2}{R^3}\right)^{1/3} \rho^i (\Delta x)^{-1/3}. \tag{S25}$$

##### D. Boundary and initial conditions

In our study we explore four different boundary/initial conditions, corresponding to four different proposed sources for the colonising bacteria.

1. If the bacteria originate from the skin, this is a Dirichlet boundary condition for the base of the outside surface, $n_N^i = const.$, and a Neumann (reflecting) boundary condition for the base of the luminal surface, $m_{N+1}^i = m_{N-1}^i$.
2. If the bacteria come from the drainage bag, this is a Dirichlet boundary condition for the base of the luminal surface, $m_N^i = const.$, and a Neumann (reflecting) boundary condition for the base of the outside surface, $n_{N+1}^i = n_{N-1}^i$.
3. If there is uniform initial contamination across the outside surface, this is an initial condition $n_p^0 = const.$, with reflecting boundary conditions for both the luminal and outside surfaces, $m_{N+1}^i = m_{N-1}^i$ and $n_{N+1}^i = n_{N-1}^i$.
4. Finally, if the bladder is already contaminated before the catheter is inserted, this is an initial condition for the bladder, $\rho_0$, and again there are reflecting boundary conditions for the catheter surfaces.

#### IV. FURTHER RESULTS OF THE MODEL

##### A. Criticality of urine production rate: washout transition

As discussed in the main text, there is a washout transition within the bladder model. In this model, bacterial dynamics in the bladder are modelled by logistic growth with dilution (Eq. 2 in the main text):

$$\frac{d\rho}{dt} = r_B \rho \left(1 - \frac{\rho}{\kappa_B}\right) - k_D \rho\,.$$

Rearranging this slightly, in order to explore the effect of the dilution term, we obtain

$$\frac{d\rho}{dt} = (r_B - k_D)\rho \left(1 - \frac{r_B \rho}{(r_B - k_D)\kappa_B}\right)\,. \tag{S26}$$

Eq. S26 takes the form of a rescaled logistic growth equation:

$$\frac{d\rho}{dt} = r_{\text{eff}} \rho \left(1 - \frac{\rho}{\kappa_{\text{eff}}}\right)\,, \tag{S27}$$

where we identify an effective growth rate, $r_{\text{eff}} = r_B - k_D$, and an effective carrying capacity $\kappa_{\text{eff}} = (r_B - k_D)\kappa_B / r_B$. Therefore, by analogy with logistic growth, we must have a steady state bacterial density equal to $\kappa_{\text{eff}}$, i.e. $\kappa_B\,(1 - (k_D/r_B))$. As $k_D \to r_B$, $\kappa_{\text{eff}} \to 0$, and thus the steady state bacterial density goes to zero. The steady state density decreases with dilution rate, down to 0 at $k_D = r$, and the time taken to achieve steady state increases, since $r_{\text{eff}} = r_B - k_D$. Hence, the bacteria grow slower, and reach a lower population density, as the dilution rate increases.

##### B. Urine production rates in the population

As discussed in the main text, the urine production rate, $\lambda$, is critical for the long-term, steady-state outcome of the model. If the dilution rate, $k_D = \lambda/V$ exceeds the bacterial growth rate $r_B$, 'washout' occurs, and there cannot be be a sustained bacterial population within the urine. The mean urine production rate (for humans) is 1 mL/min (Table 1 of the main text). As discussed in the main text (assuming the residual urine volume to be 50 mL), this mean rate is close to, but below, the critical rate, $\lambda = r_B V$ (see main text Fig. 3a).

Within the population there is a distribution of urine production rates, so while the average person falls short of the critical rate, there is a subgroup of the population who exceed it. Our model predicts that these people, with urine production rates exceeding the critical rate, $\lambda > r_B V$, are not susceptible to catheter-associated bacteriuria. Through the annual *National Health and Nutrition Examination Survey*, the CDC collects health data on a random sample of the US population. This dataset includes the urine production rate and is available online for years from 2009–2020[51]. Fig. 3b of the main text shows the distribution of urine production rates in the US adult population between 2009 and 2014, overlaid with the critical urine production rate predicted by our model for catheter-associated bacteriuria (grey dashed line), assuming parameters as in main text Table 1 (for colonisation by uropathogenic *E. coli*). From this we derive an estimate of the susceptible fraction of the population (those with urine production rates less than the threshold value, $\lambda < r_B V = 1.16$ mL/min): $72.9 \pm 0.4\%$ of females, and $68.8 \pm 0.5\%$ of males. We can also calculate the mean urine production rate across the population: $1.00 \pm 1.15$ mL/min in females, and $1.09 \pm 1.12$ mL/min in males – consistent with the 1 mL/min stated in Table 1 of the main text.

Increasing fluid intake is known to be protective against urinary tract infections (UTI)[23,52–54], and several studies have linked increased fluid intake with lower rates of catheter encrustation and blockages[30,55]. Here, we proposed a mechanism through which increased fluid intake (which is correlated with urine production rate) can be directly protective against catheter-associated bacteriuria, since this rate governs a washout transition within the bladder. Applying this, we can now quantify the reduction in risk of catheter-associated bacteriuria that is associated with a (modest) increase in fluid intake (Table S1).

##### C. Bacterial surface motility can affect steady-state bacterial density in the bladder in the high urine production rate wash-out regime

In Fig. 3a of the main text, we showed that urine production rate controls the bacterial density within the bladder at long times – determining if bacteriuria will eventually occur. In contrast, for our main parameter set, the urine

| Increase in urine production rate (mL/day) | Predicted susceptibility Female (%) | Male (%) | Relative risk adjustment Female (%) | Male (%) |
|---|---|---|---|---|
| 0 | $72.9 \pm 0.4$ | $68.8 \pm 0.5$ | - | - |
| 100 | $70.4 \pm 0.4$ | $65.5 \pm 0.5$ | $-3.4 \pm 0.8$ | $-4.7 \pm 0.9$ |
| 200 | $67.3 \pm 0.4$ | $62.2 \pm 0.5$ | $-7.7 \pm 0.8$ | $-9.6 \pm 0.9$ |
| 300 | $63.6 \pm 0.5$ | $58.4 \pm 0.5$ | $-12.7 \pm 0.8$ | $-15.1 \pm 0.9$ |
| 400 | $60.1 \pm 0.5$ | $53.6 \pm 0.5$ | $-17.6 \pm 0.8$ | $-22.1 \pm 0.9$ |
| 500 | $55.7 \pm 0.5$ | $48.6 \pm 0.5$ | $-23.6 \pm 0.8$ | $-28.3 \pm 0.9$ |

Table S1: Our model predicts that increasing fluid intake decreases susceptibility to catheter-associated bacteriuria. An increase in urine production rate of 100 mL/day is equivalent to 0.07 mL/min.

production rate has only a minimal effect on the timescale of biofilm development on the luminal catheter surface - here surface diffusivity and urethral length are more significant parameters (Fig. S6a,b; Fig. S5a).

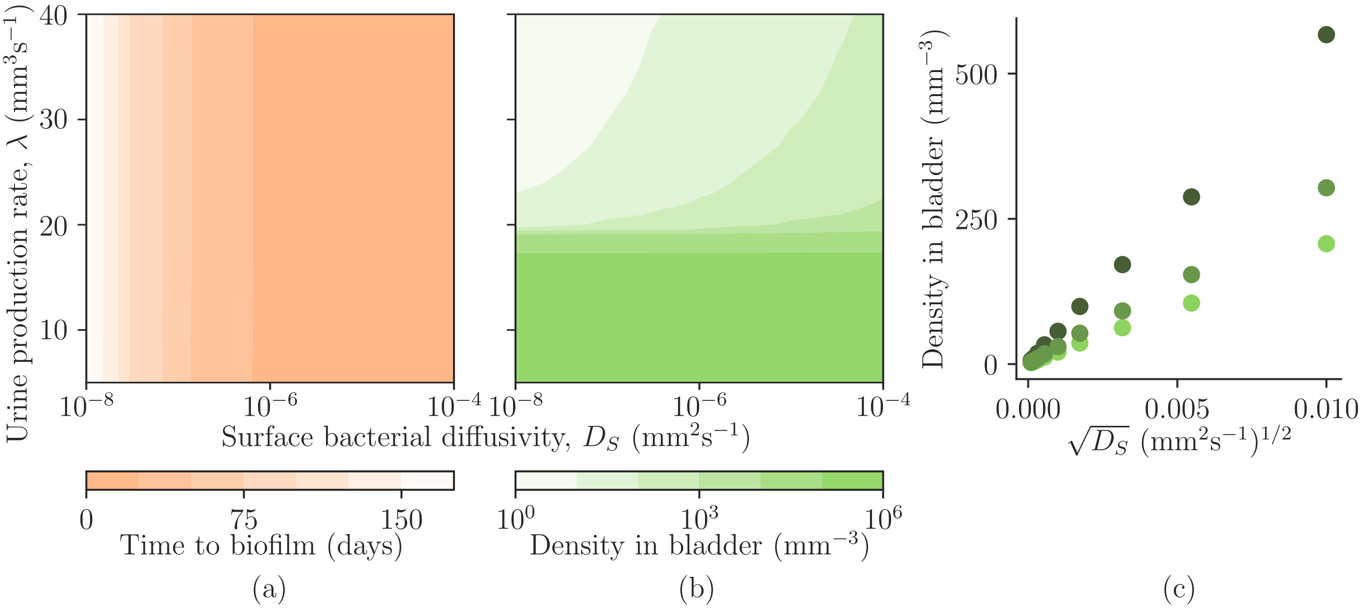


Figure S5: **Effects of varying the surface bacterial diffusivity and urine production rate.** Data is plotted for (a) the time until biofilm formation, and (b) the steady state bacterial density within the bladder. (c) Increasing the surface diffusivity leads to an increased effective coupling between the catheter surface and the bladder. This dependence is determined by the width of the FKPP wave, and scales with $\sim \sqrt{D_S}$. Plotted is the steady state bacterial density within the bladder for varying values of the bacterial surface diffusivity, for values of the urine production rate of 25 (•), 30 (•), and 35 (•) $\mathrm{mm^3s^{-1}}$. Other model parameters are as specified in main text Table 1.

Surprisingly, we find that the surface bacterial motility can also have a significant effect on the long-term outcome of the model: it alters the long-time bacterial density within the bladder in the wash-out regime where the urine production rate is high (Fig. S5b,c, $\lambda/V > r_B$). In this regime, the bacterial density within the bladder is very low, and is only sustained by the coupling with the catheter surface. Increasing the surface bacterial diffusivity effectively increases the coupling strength, increasing the bacterial density in the washed-out regime.

This effect of the surface diffusivity on the coupling between catheter and bladder is indirect. The coupling term between the surface and the bladder has no explicit dependence on the diffusivity: recall Eq. S4,

$$\frac{k_d S_c n(0,t) - k_a V_c \rho(t)}{V}.$$

However, since the steady state bacterial density on the outside catheter surface (main text Eq. 1) is effectively[1] greater than that of the bladder (main text Eq. 2), there is a 'dip' in the bacterial surface density $n(x,t)$ at the

[1] The surface density can be compared to the volume density though considering the absolute number of bacteria 'in contact' through

catheter tip (e.g. see main text Figure 5c upper). The 'steepness' of this dip is determined by the width of the FKPP wave, which is $w = 8\sqrt{D_S/r_S}$. Thus, increasing the surface diffusivity effectively increases the area of catheter that is in 'contact' with the bladder, and so increases the effective coupling strength, leading to a higher bacterial density within the bladder (Fig. S5c).

#### D. Parameter regimes for the infection timescale, and their link to patient characteristics

The parameter set of Table 1 of the main text, which underpins the data presented in the main text, corresponds to a parameter regime in which the timescale of bacterial migration up the catheter is longer than the timescale over which bacteria grow to population the bladder. We term this the slow migration / fast growth parameter regime, and we expect most clinical scenarios to fall into this regime. We note, however, that different predictions are obtained for the short-term colonisation dynamics if migration is fast compared to growth in the bladder (fast migration / slow growth regime), or if the two timescales are comparable (mixed regime).

The relevant dimensionless number controlling which regime the model is in is the ratio of the two timescales, ascension and proliferation, i.e. the ratio of the characteristic time of the Fisher wave, and the characteristic time of growth within the bladder:

$$\alpha = \frac{L}{\sqrt{r_S D_S}} \Big/ \left(\frac{\ln(\kappa_{\text{eff}}/\rho_0)}{r_{\text{eff}}}\right) = \frac{(r_B - k_D)L}{\ln\left(\frac{\kappa_B}{\rho_0}(1 - \frac{k_D}{r_B})\right)\sqrt{r_S D_S}}. \tag{S28}$$

If $\alpha \gg 1$, our model is in the slow migration / fast growth regime, and the urethral length and bacterial surface motility determine the infection timescale. If $\alpha \ll 1$, the model is in the fast migration / slow growth regime, and the bacterial growth rate in the urine and the urine production rate determine the infection timescale. The mixed regime occurs for $\alpha \approx 1$, in which case all of the above properties contribute to the infection timescale. This is the regime that is discussed in the main text (for the parameter values in Table 1 of the main text and setting $\rho_0 = 1$ since growth in the bladder occurs as soon as a single bacterium is present, we obtain $\alpha \approx 1$).

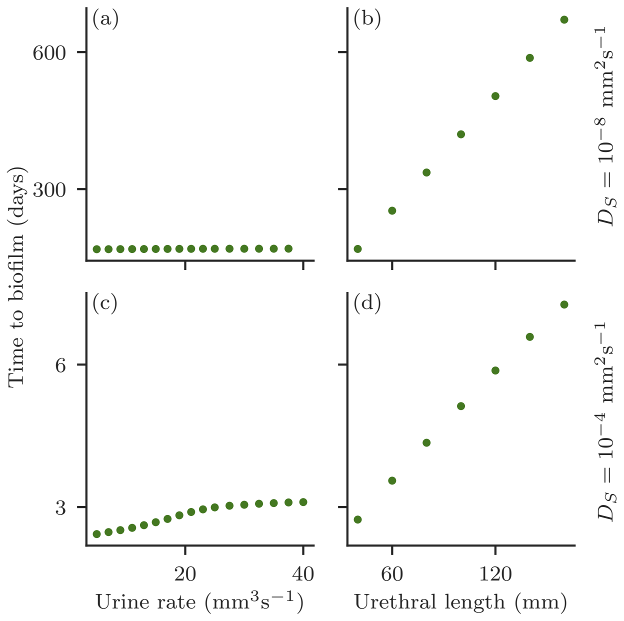


Figure S6: **The interplay of physical parameters and bacterial characteristics on the timescale of biofilm formation**. We plot the time taken for the maximum bacterial surface density on the catheter lumen to exceed $10^7$ mm$^{-2}$. Left, (a) and (c), varying the urine production rate. Right, (b) and (d), varying the urethral length. Upper, (a) and (b), have a bacterial surface diffusivity of $10^{-8}$ mm$^2$s$^{-1}$. Lower, (c) and (d), have a bacterial surface diffusivity of $10^{-4}$ mm$^2$s$^{-1}$. All other parameters are as given in main text Table 1.

---

the coupling term.

Catheter blockage is a frequent and serious complication of catheter use. Most commonly, blockage is caused when biofilms of *Proteus mirabilis* form on the catheter surface and increase the urine pH, causing crystals to precipitate[27,56]. However, non-crystalline biofilms of other uropathogens such as *E. coli* can also disrupt urine flow[5,22]. Our model cannot directly predict catheter blockage, since we do not model biofilm-associated changes in urine flow. However, as a proxy, we predict the time until the value of the surface density of bacteria at any point on the luminal surface reaches a threshold of $10^7$ bacteria per $mm^2$ (corresponding roughly to a monolayer), which we term 'time to biofilm'.

In our model, the time to biofilm formation is closely associated with the time to infection. Fig. S6 explores the effect of increasing the bacterial surface diffusivity on the timescale of infection (as measured by time to achieve a threshold surface density of bacteria). High surface diffusivity implies fast migration of bacteria up the catheter. Fig. S6a,b ($D_S = 10^{-8}$ mm$^2$s$^{-1}$, $\alpha \approx 100$) shows results corresponding to the slow migration / fast growth regime: here the timescale is controlled by migration up the catheter and the urine production rate does not play a role; equivalent results are shown also in Fig. S5a. In contrast, for the parameter set of Fig. S6c,d (with $D_S = 10^{-4}$ mm$^2$s$^{-1}$, $\alpha \approx 1$), we are well within the mixed regime and indeed the model predicts that both the urine production rate and the urethral length play a role in the timescale of infection. Notably, the bacterial surface motility is the model parameter which is least well-defined in the literature. In particular, surface motility is highly dependent on the surface 'wetness' (see Table 1 in the main text, and the supplementary background I.G. 'Motility of *E. coli*').

##### E. Comparing bacterial density patterns with experimental data

Zhang et al.[57] investigated the efficacy of a silver-PTFE coated catheter in preventing *E. coli* ascension up the catheter in an *in vitro* bladder model, in which a catheter was embedded in an agar 'urethra' with an artificial urine growth medium flowing down the catheter from a 'bladder'. The authors measured the biofilm density (via UV absorbance after crystal violet staining) on catheter sections at different positions along the catheter, to obtain bacterial density profiles, similar to those we numerically simulated in main text Figure 5, as well as measuring the time from inoculation until 'bacteriuria' (here defined as $\geq 10^3$ CFU/mL within the 'bladder').

In a separate paper arising from the same *in vitro* study, Wang et al.[58] compared bacterial ascension for a bacterial density at the 'skin' ($x = 13$ cm) of either $10^6$ cells/mL, or $10^2$ cells/mL. The shapes of the observed bacterial density distributions are strikingly similar to FKPP density wave profiles. Wang et al.[58] determined the time taken to 'bacteriuria' to be 1.8 days (silicone) or 4 days (silver-PTFE), for a 'skin' density of $10^6$ cells/mL, and 6 days (silicone) or 41 days (silver-PTFE), for a 'skin' density of $10^2$ cells/mL. Assuming FKPP dynamics (as in our model), we can estimate both $r_S$ and $D_S$ for silver-PTFE catheters, compared to silicone catheters. Our model predicts that the time taken to bacteriuria is the timescale for the asymptotic (FKPP) wavefront to reach the top of the catheter, plus the timescale over which the bacterial density grows to the detection threshold, once the bladder is contaminated. Evaluating this we find

$$\begin{aligned} r_{S;\text{Ag-PTFE}} &= 0.062 \text{ day}^{-1} & D_{S;\text{Ag-PTFE}} &= 4200 \text{ mm}^2\text{day}^{-1} \\ r_{S;\text{silicone}} &= 0.55 \text{ day}^{-1} & D_{S;\text{silicone}} &= 2400 \text{ mm}^2\text{day}^{-1} \end{aligned} \tag{S29}$$

Comparing the parameters obtained in Eq. S29 with the parameters we assumed in main text Table 1 ($r_S = 0.69$ hr$^{-1}$; $D_S = 10^{-4}$ mm$^2$s$^{-1}$), we see that our calculation (Eq. S29) results in values of $D_S \sim 10^{-2}$ mm$^2$s$^{-1}$, and $r_S = 0.02$ hr$^{-1}$. These describe a bacterial strain that is highly motile[2] and unusually slow growing. The bacterial strain used by Wang et al.[58] was a uropathogenic *E. coli* clinical isolate that has not been well characterised, and Wang et al.[58] did not report the growth rate of the bacterial strain on the artificial urine medium they utilised within the study. It is plausible that the artificial urine medium constrained the bacterial growth rate.

##### F. Global sensitivity analysis

A global sensitivity analysis was performed to identify the key parameters for two outcomes: the steady state bacterial density within the bladder, and the time taken to attain steady state within the bladder. The results of this analysis are shown in Figure S7, and summarized in main text Table 2. The key parameters controlling the long-term

---

[2] *E. coli* is known to exhibit swarming behaviours, with similar motility to as observed by Wang et al.[58], this has been observed particularly for *E. coli* cells confined between agar and a surface[35] – exactly as was the case in the setup of Wang and Zhang et al.[57,58].

outcome (here the bladder steady state density) are different to the parameters controlling the dynamics (here the time to bladder steady state).

The parameter space was sampled using SALib sample_sobol with a Sobol sample number of 16384 (corresponding to 360448 model runs), and Sobol indices calculated with SALib analyze_sobol. Bounds and distributions used can be found in Table S2.

| Parameter | | Lower bound | Upper bound | Distribution |
|---|---|---|---|---|
| Urethral length | $L$ | 40 mm | 160 mm | Uniform |
| Residual urine volume | $V$ | 10 mL | 100 mL | Uniform |
| Urine production rate | $\lambda$ | 0.3 mL/min | 3 mL/min | Uniform |
| Catheter internal radius | $R$ | 0.5 mm | 2 mm | Uniform |
| Bacterial surface diffusivity | $D_S$ | $10^{-8}$ mm$^2$s$^{-1}$ | $10^{-3}$ mm$^2$s$^{-1}$ | Log-uniform |
| Bacterial bulk diffusivity | $D_B$ | $10^{-5}$ mm$^2$s$^{-1}$ | $10^{-3}$ mm$^2$s$^{-1}$ | Log-uniform |
| Catheter surface bacterial growth rate | $r_S$ | 0.0036 hr$^{-1}$ | 3.6 hr$^{-1}$ | Log-uniform |
| Bacterial growth rate in bladder | $r_B$ | 0.0036 hr$^{-1}$ | 3.6 hr$^{-1}$ | Log-uniform |
| Catheter surface carrying capacity | $\kappa_S$ | $10^5$ mm$^{-2}$ | $10^8$ mm$^{-2}$ | Log-uniform |
| Bladder carrying capacity | $\kappa_B$ | $10^4$ mm$^{-2}$ | $10^7$ mm$^{-2}$ | Log-uniform |

Table S2: **Bounds and distributions for calculating Sobol indices.**

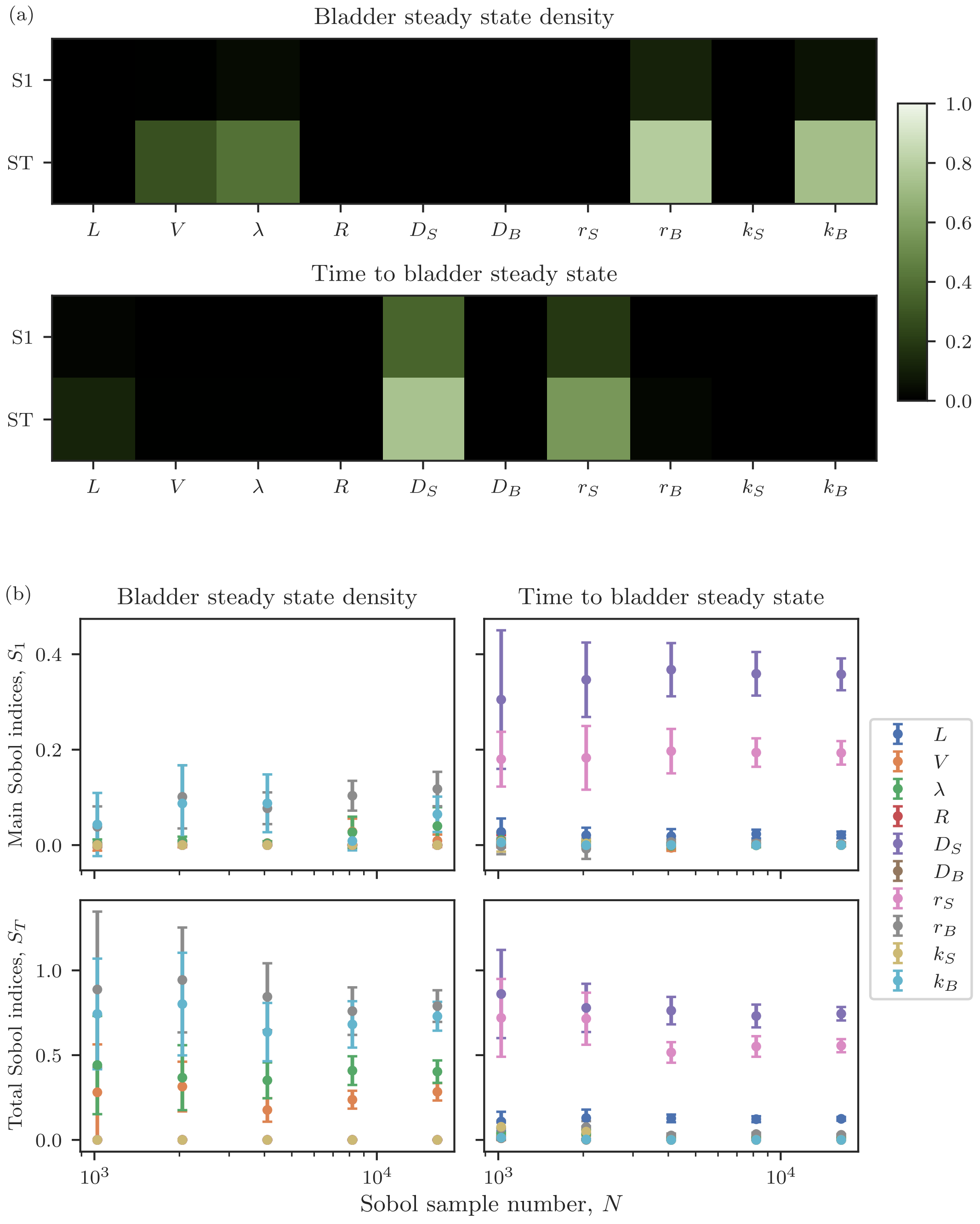


Figure S7: **Different parameters control outcomes and timescales.** (a) Main and total Sobol indices for the time to attain the steady state bacterial density in the bladder, and the value of the bacterial density in the bladder, for the parameters in Table S2. Sobol indices calculated using SALib with a Sobol sample number of 16384 (corresponding to 360448 model runs). (b) Convergence of main and total Sobol indices with increasing sample number. Plotted are the Sobol indices for each of the parameters in Table S2, with 95% confidence interval.

[35] D. B. Kearns, Nat. Rev. **8**, 634 (2010).
[36] A. Be'er and G. Ariel, Movement Ecology **7** (2019).
[37] M. Ramstedt, I. A. Ribeiro, H. Bujdakova, F. J. Mergulhão, L. Jordao, P. Thomsen, M. Alm, M. Burmølle, T. Vladkova, F. Can, et al., Macromol. Biosci. **19**, 1800384 (2019).
[38] J. Henrichsen, Bacteriol. Rev. **36**, 478 (1972).
[39] A. Gelimson, K. Zhao, C. K. Lee, W. T. Kranz, G. C. L. Wong, and R. Golestanian, Phys Rev Lett **117**, 178102 (2016).
[40] S. Spagnolie and E. Laura, J Fluid Mech **700**, 105 (2012).
[41] J. Hill, O. Kalkanci, J. L. McMurry, and H. Koser, Phys Rev Lett **98**, 068101 (2007).
[42] T. Kaya and H. Koser, Biophys J **102**, 1514 (2012).
[43] J. N. Walker, A. L. Flores-Mireles, C. L. Pinkner, H. L. Schreiber, M. S. Joens, A. M. Park, A. M. Potretzke, T. M. Bauman, J. S. Pinkner, J. A. Fitzpatrick, et al., Proc. Natl. Acad. Sci. USA **114**, E8721 (2017).
[44] S. Vogel, *Life in Moving Fluids* (Princeton University Press, Princeton, 1994), 2nd ed.
[45] B. A. Inman, W. Etienne, R. Rubin, R. A. Owusu, T. R. Oliveira, D. B. Rodriques, P. F. Maccarini, P. R. Stauffer, A. Mashal, and M. W. Dewhirst, Int J Hyperthermia **29**, 206 (2013).
[46] M. Von Smoluchowski, Z Phys Chem **92**, 129 (1917).
[47] V. G. Levich, *Physicochemical Hydrodynamics* (Prentice-Hall, Englewood Cliffs, N.J., 1962).
[48] H. J. Busscher and H. C. Van Der Mei, Clin Microbiol Rev **19**, 127 (2006).
[49] K. M. Agbavon, A. R. Appadu, and M. Khumalo, Adv Differ Equ **2019** (2019).
[50] J. D. Murray, *Mathematical Biology: I: An Introduction* (Springer-Verlag, New York, 2002), 3rd ed.
[51] Centers for Disease Control and Prevention (CDC), National Center for Health Statistics (NCHS), *National Health and Nutrition Examination Survey Data* (2009–2014).
[52] T. M. Hooton, M. Vecchio, A. Iroz, I. Tack, Q. Dornic, I. Seksek, and Y. Lotan, JAMA Intern Med **178**, 1509 (2018).
[53] Y. Lotan, M. Daudon, F. Bruyère, G. Talaska, G. Strippoli, R. J. Johnson, and I. Tack, Curr Opin Nephrol Hypertens **22** (2013).
[54] A. M. Scott, J. Clark, C. D. Mar, and P. Glasziou, Br J Gen Pract **70**, E200 (2020).
[55] R. G. Burr and I. M. Nuseibeh, Spinal Cord **35**, 521 (1997).
[56] S. A. Wilks, M. J. Fader, and C. W. Keevil, PLoS ONE **10**, e0141711 (2015).
[57] S. Zhang, L. Wang, X. Liang, J. Vorstius, R. Keatch, G. Corner, G. Nabi, F. Davidson, G. M. Gadd, and Q. Zhao, Biomat. Sci. Eng. **5**, 2804 (2019).
[58] L. Wang, S. Zhang, R. Keatch, G. Corner, G. Nabi, S. Murdoch, F. Davidson, and Q. Zhao, J. Hosp. Infect. **103**, 55 (2019).
